# The catalytic activity of microRNA Argonautes plays a modest role in microRNA star strand destabilization in *C. elegans*

**DOI:** 10.1101/2023.01.19.524782

**Authors:** Kasuen Kotagama, Acadia L. Grimme, Leah Braviner, Bing Yang, Rima M. Sakhawala, Guoyun Yu, Lars Kristian Benner, Leemor Joshua-Tor, Katherine McJunkin

## Abstract

Many Argonaute proteins can cleave RNA (“slicing”) as part of the microRNA-induced silencing complex (miRISC), even though miRNA-mediated target repression is generally independent of target cleavage. Here we use genome editing in *C. elegans* to examine the role of miRNA-guided slicing in organismal development. In contrast to previous work, slicing-inactivating mutations did not interfere with normal development when introduced by CRISPR. We find that unwinding and decay of miRNA star strands is weakly defective in the absence of slicing, with the largest effect observed in embryos. Argonaute-Like Gene 2 (ALG-2) is more dependent on slicing for unwinding than ALG-1. The miRNAs that displayed the greatest (albeit minor) dependence on slicing for unwinding tend to form stable duplexes with their star strand, and in some cases, lowering duplex stability alleviates dependence on slicing. Gene expression changes were consistent with negligible to moderate loss of function for miRNA guides whose star strand was upregulated, suggesting a reduced proportion of mature miRISC in slicing mutants. While a few miRNA guide strands are reduced in the mutant background, the basis of this is unclear since changes were not dependent on EBAX-1, a factor in the Target-Directed miRNA Degradation (TDMD) pathway. Overall, this work defines a role for miRNA Argonaute slicing in star strand decay; future work should examine whether this role could have contributed to the selection pressure to conserve catalytic activity of miRNA Argonautes across the metazoan phylogeny.

## Introduction

Multiple classes of small RNAs bind to Argonaute proteins to carry out diverse regulatory functions such as post-transcriptional gene regulation, transcriptional regulation, and programmed DNA elimination (1–4). The PIWI domain of Argonaute resembles RNase H and in many Argonautes retains the ability to cleave RNA (“slicing”) (4–14). While the role for Argonaute-mediated slicing is well-defined in some small RNA pathways, the role of slicing in the animal microRNA (miRNA) pathway is less clear.

Catalytic activity requires a tetrad of conserved residues (DEDH) (8, 9, 15, 16), with other parts of the protein also contributing supporting roles to catalysis (17–21). The catalytic or “slicing” activity of Argonaute proteins can play a role at three phases of the RNA-induced silencing complex (RISC) life cycle: small RNA biogenesis, the unwinding step of RISC assembly, and target RNA cleavage by mature RISC.

The role for slicing in small RNA biogenesis is apparent in multiple small RNA pathways in animals. Slicing by Aubergine (Aub) and Argonaute-3 (Ago3) is required for PIWI-interacting RNA (piRNA) biogenesis in flies (22, 23), and slicing activity of Mili is required for piRNA biogenesis in mice (24, 25). Notably, *C. elegans* piRNAs (21U RNAs) do not require PIWI slicing activity and instead undergo 5’ and 3’ trimming of short precursors and subsequent methylation (26–42). In another germline small RNA class, the slicing activity of CSR-1 promotes the RNA-dependent RNA polymerase (RdRP)-dependent biogenesis of CSR-1-bound 22G RNAs (43). Slicing is dispensable for biogenesis of many other classes of small RNAs, including most miRNAs (see below) and many other classes generated by RdRP-dependent modules (1).

In RISC assembly, a small RNA duplex is first loaded into Argonaute. This is followed by unwinding, in which the “passenger” or “star” strand of the duplex dissociates from the complex, leaving the single-stranded “guide” bound to Argonaute as mature RISC. Slicing of the star strand can promote unwinding (12, 44–47). The extent to which slicing of the star strand occurs and contributes to unwinding depends on the extent of base-pairing of the small RNA duplex (12, 48–50). Thus, star strands of fully complementary siRNA duplexes are efficiently cleaved (12, 44–47). In contrast, star strands of bulged miRNA duplexes are generally inefficiently cleaved, and their dissociation is usually slicing-independent (13, 51).

In mature RISC, complementary targets occupy the same position as the small RNA star strand and can likewise be cleaved by Argonaute according to similar parameters. RISCs containing the Piwi proteins cleave complementary transposon-derived and genic transcripts in some organisms, while cleavage is dispensable in *C. elegans* (22, 52–60). In *C. elegans*, target cleavage by the Argonaute CSR-1 is crucial for regulation of germline and maternal mRNA targets (6, 61–68). High complementarity generally promotes target cleavage by slicing-competent Argonautes, but the degree of complementarity required and the efficiency of slicing is guide- and Argonaute-dependent and incompletely understood (12, 13, 69–73).

In contrast to the clear role for Argonaute-mediated slicing in small RNA biogenesis, RISC assembly, and target cleavage in some animal small RNA pathways, the necessity for slicing in the miRNA pathway is less clear. The catalytic tetrad and slicing activity are conserved in many Argonautes in the miRNA pathway, despite the fact that biogenesis, unwinding, and target repression are slicing-independent for the vast majority of miRNAs. What biological roles does slicing play in the miRNA pathway that maintain selective pressure on the critical catalytic residues?

Biogenesis of animal miRNAs is generally carried out by sequential Microprocessor- and Dicer-mediated cleavage steps (5, 74–76), and is generally independent of Argonaute-mediated slicing, with at least one clear exception: miR-451 is generated in a Dicer-independent manner that requires cleavage of its unusual precursor hairpin by Argonaute (77–79). In early *C. elegans* work, elevated miRNA precursors were observed upon RNAi of the miRNA Argonautes (<u>A</u>rgonaute-like <u>g</u>ene, ALG-1 and -2) (5), and rescue of ALG mutants with catalytic dead overexpression transgenes resulted in upregulation of partially cleaved precursors (13). These results initially suggested a role for ALG-1/2 in general miRNA biogenesis, but this interpretation should be revisited given our current understanding of miRNA biogenesis and more elegant genetic tools (i.e. CRISPR).

Unwinding of animal miRNAs is also largely slicing-independent and is aided by a wedging function of Argonaute’s N domain and residues in the PIWI domain (51, 80, 81). One exception to this is the miR-486 guide:star duplex which is completely complementary (like an siRNA) and requires Argonaute to cleave the star strand for unwinding and activation of miR-486-RISC (82). Interestingly both slicing-dependent miRNAs are required for erythropoiesis, and rescuing expression of both miR-451 and miR-486 bypasses the requirement for Ago2-mediated slicing in erythropoiesis (82). Nonetheless, miR-451 and miR-486 are not conserved outside of vertebrates (83), so these described roles for slicing do not explain why miRNA Argonautes would retain their catalytic activity in other animals.

What about target cleavage? In animals, miRNAs generally bind their targets with incomplete complementarity (84–87), which is sufficient for repression of the target through translational repression, deadenylation, and/or decay (88–93), and the vast majority of endogenous miRNA targets are repressed in a GW182-dependent manner that does not require target cleavage, with a few known exceptions (94–104). Because miRISC can efficiently promote target repression in the absence of slicing, slicing may be completely dispensable for target repression, even though it does occur in a limited number of miRNA:target interactions (98–100, 102). One complication to answering this question in mammals is the multiple small RNA populations bound by the slicing-competent miRNA Argonaute Ago2. Outside of the vertebrate hematopoietic system, catalytic activity of Ago2 is also required for trophectoderm development, a function that is presumed to be due to Ago2’s role in binding endogenous siRNAs (105); Ago2-mediated slicing plays additional as yet unidentified essential roles in intra-embryonic tissues outside the hematopoietic lineage (77, 82). In other animals – such as in *C. elegans* and *Drosophila melanogaster* – endo-siRNAs, exo-siRNAs, and miRNAs are bound by distinct Argonaute proteins (14, 106–114), so conservation of siRNA function would not exert selective pressure on the miRNA Argonautes. The dedicated roles of Argonaute proteins in these systems also makes these models excellent contexts for assessing the roles of slicing in the miRNA pathway.

A study by Bouasker and Simard previously examined the role of slicing in the miRNA pathway in *C. elegans* (13). By rescuing null mutations and RNAi of the miRNA Argonautes with overexpression of catalytically-inactivated Argonautes, Bouasker and Simard observed strong phenotypic effects, suggesting that catalytic activity in the miRNA pathway is important to animal physiology (13). Although small RNAs and mRNAs were not profiled globally, northern blots of select miRNAs showed incomplete precursor cleavage of some miRNAs in the context of overexpression of catalytic-dead Argonaute, suggesting a role for slicing in miRNA biogenesis (5, 13).

We set out to revisit the role of the catalytic tetrad in *C. elegans* miRNA Argonautes using CRISPR/Cas9-mediated genome editing and deep sequencing of small RNAs and mRNAs. Here we show that slicing plays a role in star strand ejection and decay, and that this is more prominent in the Argonaute ALG-2 than ALG-1. However, the observed effects are only modest when using genome editing to inactivate slicing. A very mild impact on target regulation was observed by RNA-seq. Consistent with this, animal physiology was like wild type. Thus, revisiting this question with state-of-the-art techniques shows that the dramatic effects observed in the previous study may have been due to overexpression (as necessitated by transgenic technologies at that time). This work demonstrates a modest role for miRNA-guided RNA cleavage in an invertebrate organism. Selective pressure to retain catalytic activity of miRNA Argonautes may be subtle or only apparent in harsher non-laboratory “real world” conditions.

## Methods

### C. elegans culture

Nematode culture was performed on NGM plates seeded with OP50 (115). Worms were maintained at 20°C except where otherwise noted. Embryo samples were harvested by hypochlorite treatment. For L4 and young adult samples, L1s were hatched overnight in M9 supplemented with cholesterol, then grown on NGM with OP50 for 48h at 20°C (for L4s) or for 44h at 25°C (for young adults).

### Genome editing

Guide RNAs were designed as close to the site of the edit as possible, while also favoring high GC content and favorable CRISPRscan scores (116, 117). Alt-R crRNAs (IDT) were pre-annealed with tracrRNA at 10μM by heating to 95°C for 5 minutes in IDT duplex buffer, then cooling to room temperature. Annealed guide RNAs were then assembled with Cas9 (IDT) at 2μM Cas9 and 4μM of total guide RNAs. Injection mixes also included donor molecules with 35-nt homology arms for homology-directed repair (118); donor molecules were 100-150ng/µl (since all donors were relatively short). As a marker of CRISPR efficiency, a guide RNA and donor for *dpy-10* was included in all injection mixes (119).

For *mir-63* and *mir-72* mutations, two rounds of CRISPR were performed. First, the *mir-63* and *mir-72* precursors were deleted and replaced by a cleavage site for an efficient crRNA (120) either in the wild type or the *alg-1(AEDH); alg-2(AEDH)* background (strains MCJ488 and MCJ511). In these backgrounds, CRISPR was performed again to reintroduce the mutant versions of the *mir-63* and *mir-72* precursors.

For 3xFLAG-tagged ALG-2, two rounds of CRISPR were performed. The first generated a deletion near the N terminus of ALG-2 harboring a cleavage site for an efficient crRNA (*cdb12)*. This allele was backcrossed twice to N2, then introduced into a background stably expressing Cas9 (oxSi1091); a second round of CRISPR was performed in this strain by injecting crRNA at 1µM with donor DNA (∼100ng/µl) to restore all deleted sequence while also introducing the 3xFLAG and a short linker sequence in frame (*cdb51)*. The resulting locus was then backcrossed twice to N2. To generate the AEDH mutations in the context of 3xFLAG-tagged ALG-1/2, the 3xFLAG-tagged strains of ALG-2 (MCJ133) or ALG-1 (QK67, a gift from John Kim) were injected with CRISPR reagents targeting the first residue of the catalytic tetrad. Strains, alleles and oligos used in this study are listed in Supplemental Tables S1 and S2.

### Western blotting

Samples were resuspended by matching worm pellet volume with equal volume 2x lysis buffer and supplementing with additional 1x lysis buffer (30mM HEPES pH=7.4, 50mM KCl, 0.2% Triton X-100, 2mM MgCl_2_, 10% glycerol, 2mM DTT, 1x cOmplete, Mini, EDTA-free Protease Inhibitor Cocktail, 1x PhosSTOP phosphatase inhibitor tablets). Samples were lysed by 10 cycles (30s on, 30s off) of sonication in a Bioruptor Pico, then clarified by centrifugation at >15,000*g* for 20 minutes at 4°C. Samples were quantified by Pierce Reducing Agent Compatible BCA Protein Assay and run on 4-20% TGX gels (Biorad). Membranes were probed with anti-FLAG M2 antibody (Sigma) diluted 1:1000, anti-tubulin DM1A diluted 1:1000 (gift from Kevin O’Connell), anti-H3 1B1B2 diluted 1:1000 (abcam); all hybridizations were carried out in 5% milk in Tris buffered saline with .1% Tween-20.

### Brood size assays

Brood size assays were conducted at 25°C after maintaining the strains at 25°C for one generation before the assay. Specifically, MCJ275, MCJ277, MCJ288, N2, RF54, and WM53 strains were grown for ∼96hr at 25°C, after which L4 worms were singled onto individual plates. Worms were allowed to grow and lay eggs for 24h and moved to new plates until the worms stopped laying embryos. Plates with embryos were incubated for another 24h before larvae and unhatched embryos were counted.

### RNAi assays

RNAi plasmids targeting *alg-1* (MSPO171), *alg-2* (MSPO163) and the RNAi control plasmid L4440 were gifted by Martin Simard (13). An RNAi clone targeting the 3’UTR of *alg-2* was gifted by Amy Pasquinelli (121). RNAi plates were prepared with 1µg/mL IPTG + 1µg/mL Carbenicillin and stored at 4°C for less than two weeks prior to use. Synchronized L1 populations of worms were generated by alkaline hypochlorite lysis of gravid adults followed by hatching overnight in M9 with 1mM cholesterol at 20°C. (Note that the *gpap-1* SNP is absent from all strains scored in this assay; see also Table S1.) Approximately 100 L1 worms were plated on RNAi food or vector control (L4440) and incubated at 15°C for 96h, after which they were scored as fertile adults, dead, slow growing, sterile or males as in (13).

### Embryonic lethality assays

Synchronized L1 populations of MCJ275 and N2 worms were generated as described above. L1s were then plated on NGM plates with OP50. Worms were grown at 15°C for 96h and fertile adults were singled to individual plates and allowed to lay eggs overnight. The following day the adult worms were removed, and the eggs incubated overnight at 15°C. Larvae and unhatched embryos were counted the next day to evaluate embryonic lethality.

### Quantitative miRNA-Taqman

For each sample, 9.96ng of total RNA was used to perform 5μL reverse transcription reactions using the TaqMan MicroRNA Reverse Transcription kit (ThermoFisher). After RT, 1.33μL of 1:4-diluted RT reaction was used in 5μL qPCR reactions with Taqman miRNA probes and Taqman Universal Mastermix II with UNG (ThermoFisher). Technical triplicates were run for each reaction on the Applied Biosystems QuantStudio Pro 6. Standard curves were generated by similarly performing miRNA-Taqman on a dilution series of an RNA oligo of the sequence of interest (Horizon Discoveries).

### Deep sequencing library construction and data analysis

Samples were snap frozen, then resuspended in Trizol and vortexed for 15 minutes at room temperature. Thereafter, total RNA was purified according to the Trizol manufacturer’s instructions (ThermoFisher).

For preparing small RNA libraries, RNA oligos listed in Table S2 were spiked into total RNA at a concentration of 1pg/µl in 100ng/µl total RNA. Small RNA libraries were prepared using the NEBNext Small RNA Library Prep kit (NEB), with some modifications. Size selection was only performed after reverse transcription, using 8% urea gels to purify 65-75nt products. RT products were treated with RNAse H for 30 min at 37°C prior to gel purification. After excision, products were eluted overnight in 700µl 0.3M NaCl in TE. Products were then purified by isopropanol precipitation, washed in 80% ethanol, and resuspended in 40µl 50mM Tris-HCl 75mM KCl 3mM MgCl2 pH 8.3 before proceeding with the NEBNext protocol. Samples were sequenced on a HiSeq 3000 or NextSeq 550 (Illumina). Raw data can be accessed under SRA Bioproject accession number PRJNA922944.

Small RNA libraries were trimmed using Cutadapt 3.4 (122), then mapped using bowtie2 2.4.4 with settings “--no-unal --end-to-end –sensitive” (123). BAM files were sorted and indexed using samtools 1.13, then assigned to miRNAs using htseq 0.13.5 (124, 125). For genome mapping and miRNA assignment, custom reference files were used which included the spiked in RNA oligos. Statistical analysis was performed using DESeq2 and GraphPad Prism (126). For spike-in normalization (Figure S3), size factors for DESeq2 analysis were calculated by dividing spike-in reads in each sample by the median number of spike-in reads per sample. For all other analyses, default DESeq2 normalization was performed using all endogenous miRNA-mapping reads (spike-in reads were excluded from analysis).

For RNAseq, libraries were prepared by Psomagen using the Truseq Stranded Total RNA Library Prep Kit and sequenced on a NovaSeq6000 S4 (Illumina). Low quality bases and adaptors were trimmed from sequence reads using Cutadapt v2.7 with parameters “–q 20 -minimum-length 25” (122). Resulting reads were mapped to WS277 using HISAT2 v2.1.0 with default parameters (127). Aligned reads were counted using subread featureCounts v1.6.4 with default parameters, except that “-s” option was used to specify appropriate strands (128). Differential expression analysis was done with DESeq2 v1.22.1 (ref4) in v3.5.1 R version (126).

To calculate predicted miRNA:star duplex structures and calculate minimum free energy (MFE), RNAcofold was used with default parameters, as part of the ViennaRNA package (version 2.4.18).

### Reagents

The following reagents were used: Alt-R crRNAs (IDT), Cas9 (IDT), Trizol (ThermoFisher), TaqMan MicroRNA Reverse Transcription kit (ThermoFisher), Taqman miRNA probes (ThermoFisher), Taqman Universal Mastermix II with UNG (ThermoFisher), NEBNext Small RNA Library Prep kit (NEB), RNA oligonucleotides (Horizon Discoveries). The following specialized instruments were used: QuantStudio Pro 6 (Applied Biosystems), HiSeq 3000 (Illumina), NextSeq 550 (Illumina), NovaSeq6000 S4 (Illumina), Bioruptor Pico (Diagenode).

### Biological Resources

All strains used in the study are listed in Table S1.

### Statistical Analyses

Statistical analysis for deep sequencing datasets was performed using DESeq2 using default settings except when spike-ins were used as noted (126). For analysis of standard data types, the Analysis function in GraphPad Prism was used to select and perform the appropriate statistical test. At least three (in most cases four) replicates of each genotype were used for all experiments.

## Data Availability/Sequence Data Resources

All deep sequencing raw data can be accessed under SRA Bioproject PRJNA922944.

## Data Availability/Novel Programs, Software, Algorithms

Not applicable.

## Web Sites/Data Base Referencing

MirGeneDB 2.1 (https://mirgenedb.org/), miRBase (https://www.mirbase.org/), WormBase (https://wormbase.org/)

## Results

### Mutation of catalytic tetrad in both miRNA Argonautes does not impact physiology

To revisit the role of miRISC-mediated slicing in invertebrates, we used CRISPR/Cas9-mediated genome editing to mutate the DEDH catalytic tetrad to AEDH in the major *C. elegans* miRNA Argonautes, ALG-1 and ALG-2 (Figure 1A). (The germline-specific miRNA Argonaute ALG-5 lacks conservation in any of the positions corresponding to DEDH and therefore likely lacks catalytic activity (Figure 1A).) These AEDH mutations did not affect protein abundance (Figure S1). This mutation has previously been shown to completely inactivate catalytic activity in orthologous Argonautes *in vitro* and *in vivo* (9, 77). Although the molecular phenotypes presented below are consistent with a slicing defect, a caveat to interpretation of this study is that we have not validated slicing activity of these variants of ALG-1 and -2 directly in slicing assays (see also Discussion).

**Figure 1.**
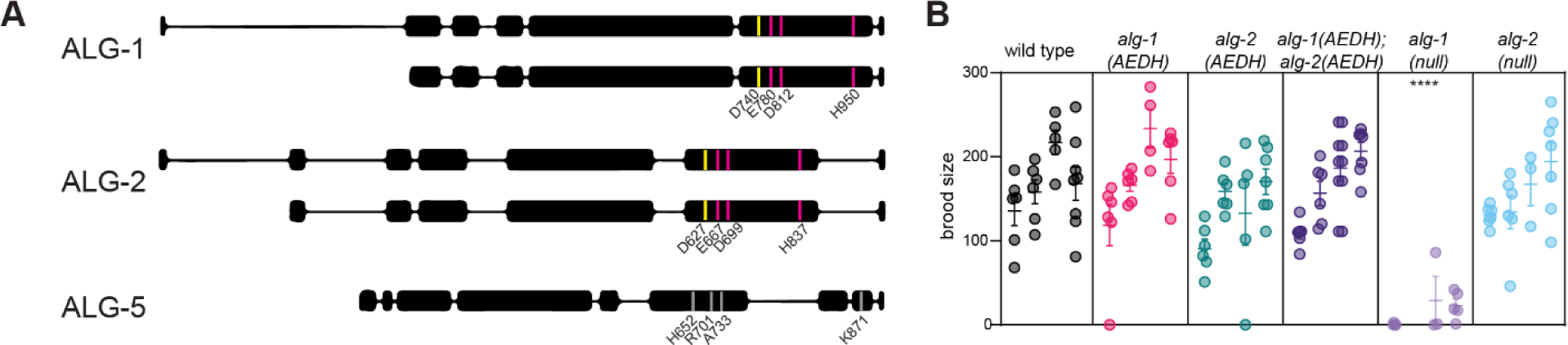
Mutation of catalytic tetrad in both miRNA Argonautes does not impact physiology. A) Schematic of miRNA Argonaute genes in *C.* with positions encoding residues of catalytic tetrad in ALG-1 and ALG-2 highlighted. ALG-5 residues in equivalent positions are not conserved. For ALG-1/2, positions in short isoform are denoted. Yellow position highlights aspartate that is mutated to alanine in “AEDH” mutants. B) Fecundity assay. Each dot represents one brood; four replicates conducted shown side by side for each genotype. (Only three replicates were done for *alg-1(null)*.) All mutants were compared to wild type by one-way ANOVA with Dunnett’s test. Only *alg-1(null)* shows significant difference from wild type. ****p<0.0001. Assay was performed at 25°C.

Mutant strains were superficially wild type. Negligible lethality was observed. Brood size assays showed that single *alg-1(AEDH)* or *alg-2(AEDH)* or double *alg-1(AEDH); alg-2(AEDH)* mutants all had wild-type like fecundity (Figure 1B). In comparison, *alg-1(null)* animals had compromised fecundity, as previously observed (5, 129). Therefore, *alg-1(AEDH)* and *alg-2(AEDH)* largely retain wild type function and do not mimic loss-of-function alleles.

These results are in contrast to a previous study in which catalytic dead versions of Argonaute were over-expressed in an Argonaute loss-of-function background (13). In that case, catalytic-dead versions of Argonaute only partially rescued viability and provided no rescue of fertility (all animals were dead or sterile). To more directly compare the phenotypes of the *alg-1(AEDH)* and *alg-2(AEDH)* CRISPR strains to the strains in the Bouasker and Simard study, we performed identical RNAi assays. In these assays, *alg-1* RNAi was performed on wild type, *alg-2(null)* and *alg-2(AEDH).* Likewise, *alg-2* RNAi was performed on wild type, *alg-1(null)* and *alg-1(AEDH).* The vector control and *alg-1* and *alg-2 CDS* RNAi clones used in Bouasker, et al. were used, as well as an additional *alg-2* RNAi clone that targets the *alg-2* 3’UTR for enhanced specificity (121). ALG-1 and -2 function largely redundantly (5); thus, as expected, RNAi of the other ALG paralog caused sterility and lethality in the *alg-1(null)* and *alg-2(null)* backgrounds (Figure S2). However, RNAi of the other ALG paralog did not induce lethal or sick phenotypes in the *alg-1(AEDH)* or *alg-2(AEDH)* backgrounds (Figure S2). Thus, consistent with the wild-type like phenotype of the *alg-1(AEDH); alg-2(AEDH)* double mutant, *alg-1(AEDH)* and *alg-2(AEDH)* do not behave as loss of function alleles in this assay. Taken together, this suggests that the previous study was impacted by expression level of the catalytic dead Argonaute and that, in fact, slicing is not required for grossly normal physiology and fitness.

### Some miRNA star strands are slightly elevated in the absence of slicing

Previous work in other models showed that the catalytic activity of Argonaute in the miRNA pathway is crucial for the biogenesis of a Dicer-independent miRNA and the proper loading of an siRNA-like miRNA (77, 78, 82). To determine the impact of catalytically inactive Argonautes on *C. elegans* miRNAs, we profiled miRNA star and guide strands from embryos, L4 stage larvae and young adults. We chose to profile these stages because of prominent roles of miRNAs in these stages and the differential expression of the miRNA Argonautes across these stages (121, 129, 130).

Exogenous RNA oligos were spiked in prior to library prep to control for potential large-scale changes in miRNA populations. Spike-in normalization (Figure S3) and normalization to all endogenous miRNA reads (Figure 2) yielded similar results, demonstrating that wholesale changes in small RNA population did not occur. Normalization to all endogenous miRNAs resulted in slightly lower variance across replicates and thus yielded slightly more significantly altered small RNAs (Figure S3 and Figure 2); thus, we proceeded with this type of normalization for all other analyses.

**Figure 2.**
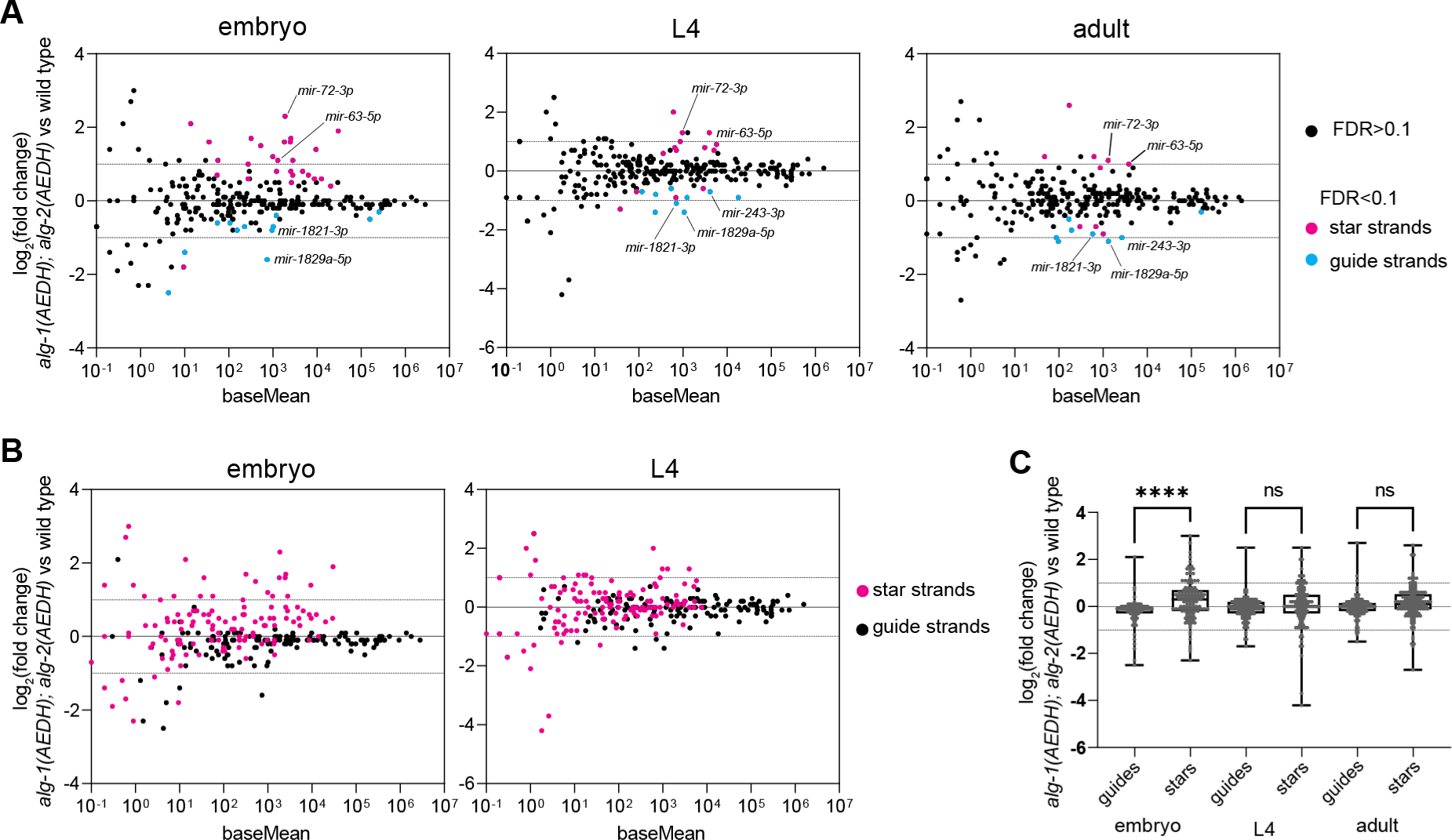
Some miRNA star strands are slightly elevated in the absence of slicing. A) MA plot showing average abundance on X-axis and log_2_(fold change) on Y-axis. miRNAs showing significant changes in *alg-1(AEDH); alg-2(AEDH)* versus wild type are shown in blue (miRNA guide strands) or pink (star strands) (DESeq2 FDR<0.1). Three (L4) or four (embryo and young adult) biological replicates of each genotype were analyzed. (B) MA plot as in (A), with all guide strands shown in black and all star strands in pink. (C) Summary of log_2_(fold change) values for all guides or star strands in each stage in *alg-1(AEDH); alg-2(AEDH)* versus wild type. Log_2_(fold change) of star strands were compared to those of guide strands in the same sample by one-way ANOVA followed by Sidak’s multiple comparisons test. ****p<0.0001

Embryo samples showed the greatest number of significantly altered miRNAs. Twenty-six miRNA strands were upregulated, all of which are miRNA star strands (Figure 2A, Tables S3-S5). Subtle star strand upregulation was also widespread such that star strands were upregulated overall compared to miRNA guide strands in embryos (Figure 2B-C). Ten and six star strands were also upregulated in L4 larvae and adults, respectively (Figure 2A, Tables S6-S7). Changes in miRNA abundance were further validated by Taqman qRT-PCR for a select set of miRNAs (Figure S4). The upregulation of star strands suggests that slicing promotes unwinding, as previously observed for siRNAs (12, 44–46). The amplitude of these changes is modest compared to the guide:star strand ratio, which is ≥10-fold for most miRNAs (131). Therefore, the putative slicing-dependent unwinding of these star strands occurs alongside a more dominant slicing-independent pathway. This low amplitude of star strand elevation suggests that only a very small proportion of miRNA guide strand function would be impaired and may not result in observable guide strand loss-of-function. Few miRNAs were also downregulated (Figure 2, Table S5-S7). These were mostly miRNA guide strands, including *mir-243-3p, mir-1821-3p,* and *mir-1829a-5p*, which were each downregulated in more than one developmental stage. Again, the amplitude of changes is modest, suggesting that slicing is not strictly required for the biogenesis of these miRNAs (Figure 2). The mechanism of the downregulation of these guide strands is not understood, though one possible model is explored below.

The contribution of slicing to star strand unwinding may predict that embryos would be more vulnerable to catalytic-dead mutations when grown at low temperature, where increased duplex stability may favor slicing-dependent unwinding (50). We tested this by growing animals at 15°C starting at L1 and scoring hatching rate of their progeny. The viability of the *alg-1(AEDH); alg-2(AEDH)* embryos (99.7%, n=364) was similar to that of wild type (99.4%, n=360). Thus, growth at mild low temperature does not reveal severe phenotypes of the *alg-1(AEDH); alg-2(AEDH)* background. Future work should examine more acute cold shock paradigms as a setting in which slicing may play a more prominent role.

### Slicing plays a greater role in ALG-2 unwinding than that of ALG-1

We sought to understand why embryo samples displayed more widespread elevation of star strands upon inactivation of slicing than other stages. We hypothesized that the larger effect in embryos might be due to a difference in relative expression of the Argonaute paralogs ALG-1 and ALG-2. ALG-2 is expressed earlier in embryogenesis than ALG-1, and its expression decreases after embryogenesis (129, 130). Furthermore, the most abundant embryonic miRNA family, *mir-35-42*, largely depends on ALG-2 expression for its stabilization (100, 129, 132). Together these data suggest that the ratio of ALG-2:ALG-1 is higher in embryos than later stages. Therefore, the larger effect of slicing deficiency in embryos may be due to a greater dependence of ALG-2 unwinding on slicing than that of ALG-1.

To test this hypothesis, we profiled small RNAs from embryos that were single mutants, with slicing inactivated in either ALG-1 or ALG-2. Single *alg-1(AEDH)* mutant embryos displayed very few significant changes in miRNA abundance (Figure 3A, Table S8). In contrast, *alg-2(AEDH)* single mutants showed nearly as many significant changes as *alg-1(AEDH); alg-2(AEDH)* double mutant embryos (Figure 3A, Table S9). Below the threshold of significance, star strands were globally slightly elevated in the *alg-2(AEDH)* single mutants, again mimicking the *alg-1(AEDH); alg-2(AEDH)* double mutant (Figure 3B). Therefore, the elevation of star strands in *alg-1(AEDH); alg-2(AEDH)* is largely due to the sensitivity of ALG-2 to catalytic inactivation. Samples harvested from later stages may show subtler changes in miRNA abundance due to a higher ratio of ALG-1:ALG-2 expression coupled with a lower sensitivity of ALG-1 to inactivation of slicing.

**Figure 3.**
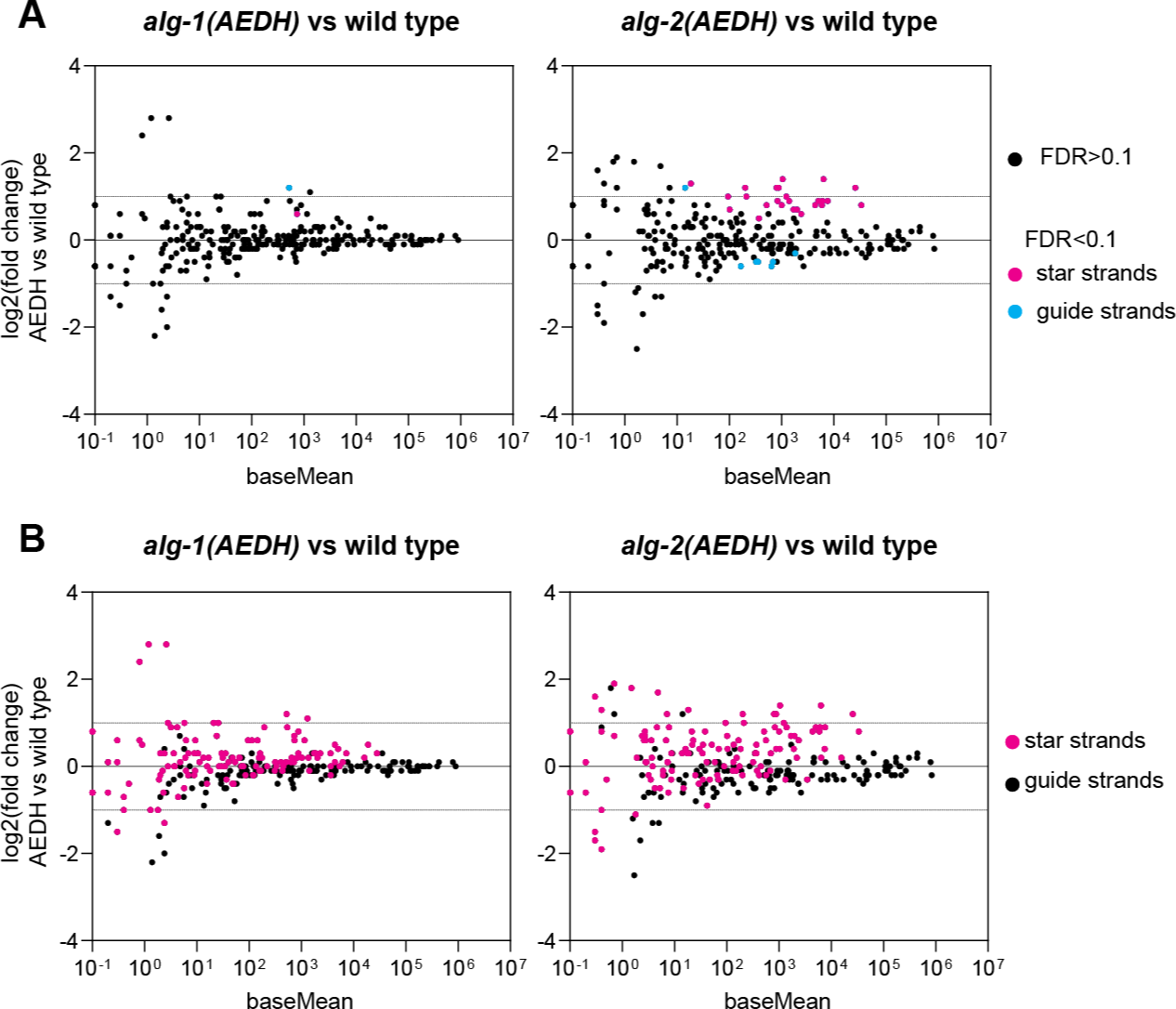
Slicing activity of ALG-2 plays a greater role in star strand destabilization than that of ALG-1. A) MA plot showing average abundance on X-axis and log_2_(fold change) on Y-axis. miRNAs showing significant changes in *alg-1(AEDH)* or *alg-2(AEDH)* versus wild type embryos are shown in blue (miRNA guide strands) or pink (star strands) (DESeq2 FDR<0.1). Four biological replicates of each genotype were analyzed. (B) MA plot as in (A), with all guide strands shown in black and all star strands in pink.

### High duplex stability favors a greater role for slicing in unwinding

While many miRNA star strands were elevated in embryos, very few were elevated in adults and L4, and these were largely consistent across both stages. We further examined the properties of these guide:star duplexes to determine if any distinctive feature might predict the extent to which a miRNA duplex depends on slicing for unwinding.

The star strands *mir-63-5p* and *mir-72-3p* were upregulated across all three stages examined. We observed that *mir-63* and *mir-72* both had relatively low minimum free energy (MFE) of the guide:star duplex compared to other miRNAs (they are both in the lowest 15% of MFEs across all miRNA duplexes) (Figure 4A-B). We also observed that the guide strands of these duplexes displayed a high rate of 3’ end trimming (within the top 12% of all guide strands) (133) (Figure 4A).

**Figure 4.**
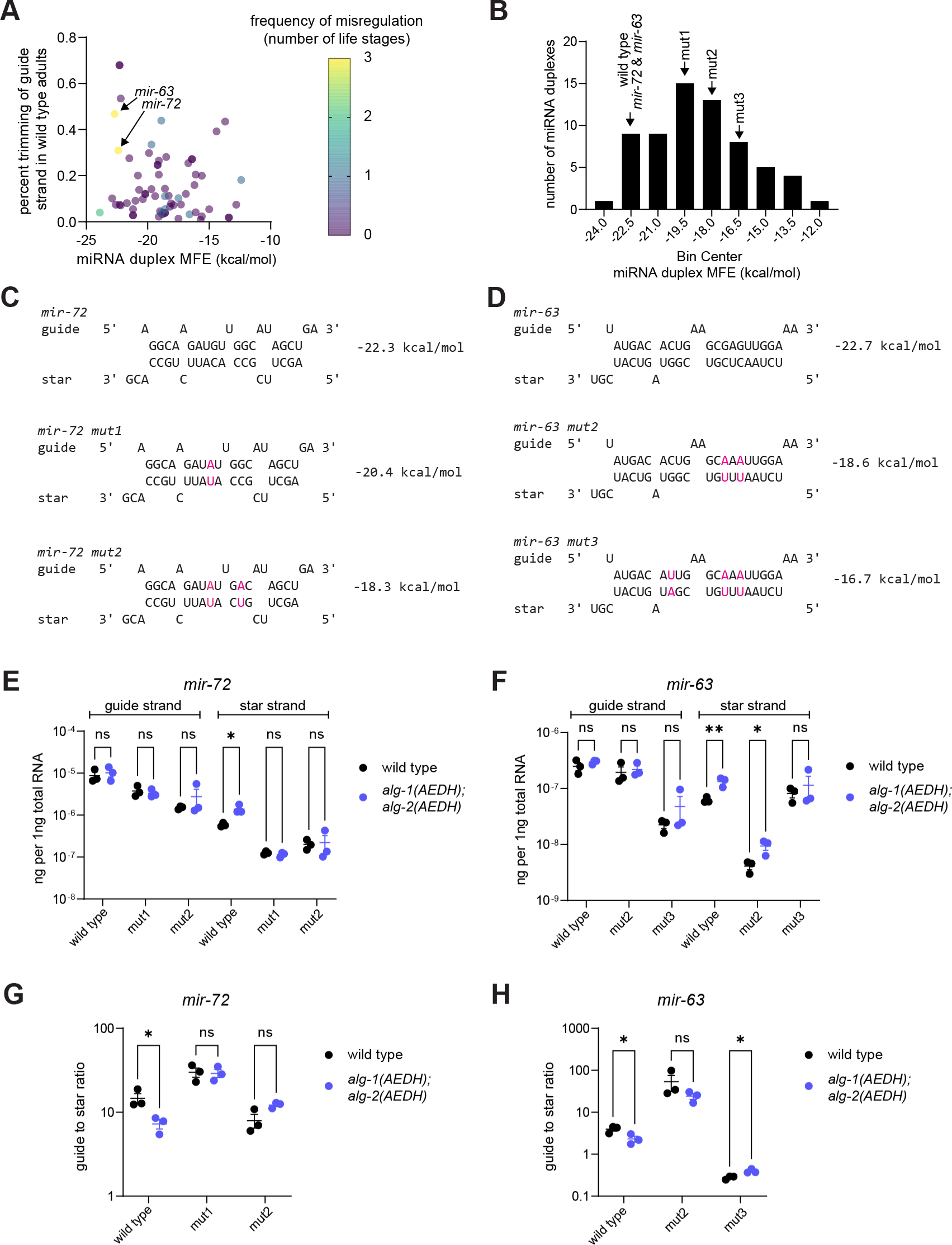
High duplex stability favors a greater role for slicing in unwinding. A) Percent trimming of guide strand and duplex stability (minimum free energy) of each miRNA duplex. Frequency of star strand misregulation in *alg-1(AEDH); alg-2(AEDH)* among three stages (embryo, L4, adult) is color-coded. B) Histogram of duplex stabilities with bins containing *mir-63* and *mir-72* and their mutant variants indicated. A-B) Only miRNAs with sufficient abundance for trimming analysis in (133) are shown. C-D) Predicted structure of *mir-63* and *mir-72* and engineered mutations in each duplex. E-F) Absolute quantification by qPCR of *mir-63* and *mir-72* guide and star strands in L4 samples (three biological replicates for each genotype). G-H) Ratio of guide to star strand calculated from absolute quantification. E-H) Unpaired t-tests, **p-value < 0.01, *p-value < 0.05.

One possible model is that the high stability of the guide:star duplex slows slicing-independent unwinding (48, 50, 134). The presence of the star strand could favor a structural conformation in which the 3’ end of the guide strand is not bound by the PAZ domain, leaving it vulnerable to exonucleolytic trimming (135, 136). Thus, a longer dwell time of the star strand could lead to higher prevalence of trimmed guide strand molecules. The slow rate of slicing-independent unwinding and thereby long dwell time of the star strand could allow the inefficient slicing-dependent star strand ejection mechanism to play a relatively larger (though still minor) role in miRISC maturation.

To test this model, we performed CRISPR to make mutations that are predicted to preserve overall base pairing structure while decreasing duplex stability (i.e. increasing MFE) (Figure 4B). These mutations consisted of one to three mutations on each strand of the miRNA duplex, replacing G:C base pairs with A:U base pairs (Figure 4C). Mutations were introduced near the middle of the duplex (avoiding the terminal four base pairs) in an attempt to preserve thermodynamic asymmetry and strand selection (137–139).

We then examined the mutant duplex miRNAs in an otherwise wild type or the *alg-1(AEDH); alg-2(AEDH)* background to determine whether decreasing duplex stability also reduces the dependence of star strand unwinding on slicing. To measure both star strand stabilization and strand selection, we performed qPCR in L4 samples calibrated with standard curves of synthetic RNA oligos to enable absolute quantification (Figure S5). For both wild type *mir-72* and *mir-63*, qPCR confirmed upregulation of the star strand in the *alg-1(AEDH); alg-2(AEDH)* mutant background with no concomitant effect on the guide strand (Figure 4E-F). For *mir-72*, either one or two mutations that reduced thermodynamic stability of the guide:star duplex also abrogated the effect of *alg-1(AEDH); alg-2(AEDH)* on the star strand abundance (Figure 4E). This suggests that the dependence of the wild type duplex on slicing for complete unwinding is due to high thermodynamic stability of the miRNA:star duplex. For both *mir-72* mutant duplexes, the guide strand selection remained high, arguing against an alternative model in which star strand effects are altered by increased star strand loading into Argonaute (Figure 4G).

For *mir-63*, two mutations in the duplex (*mir-63_mut2)* did not abolish the increased star strand abundance observed in the *alg-1(AEDH); alg-2(AEDH)* background (Figure 4F). A third mutation in the duplex of *mir-63* (*mir-63_mut3)* caused a switch in strand selection that favored loading of *mir-63-5p* over the erstwhile guide strand *mir-63-3p*, making it difficult to interpret the effects of this mutation in the *alg-1(AEDH); alg-2(AEDH)* background (Figure 4H).

Overall, the *mir-72* experiments show that duplex thermodynamic stability is one factor that influences dependence on slicing for unwinding (Figure 4E). At the same time, the weak correlation between thermodynamic stability and dependence on slicing (color and MFE - Figure 4A) and the lack of effect of two mutations in *mir-63* (Figure 4F) together show that other unknown factors also influence dependence on slicing for unwinding.

### Minor changes in star strand abundance result in subtle changes in target repression

As discussed above, physiology of the *alg-1(AEDH); alg-2(AEDH)* double mutant animals is grossly normal (Figure 1). Next, we performed RNA-seq to determine whether changes in miRNA abundance (Figure 2) or absence of target cleavage led to a more subtle molecular phenotype. Very minor changes in gene expression were observed. No genes were significantly altered in expression in L4 stage *alg-1(AEDH); alg-2(AEDH)* animals compared to wild type, whereas only one or two were altered in embryo and adult, respectively (Figure 5A, Tables S10-S12). These changes were both down-regulation of protein-coding genes (not an up-regulation that might be expected from de-repression of miRNA target genes).

**Figure 5.**
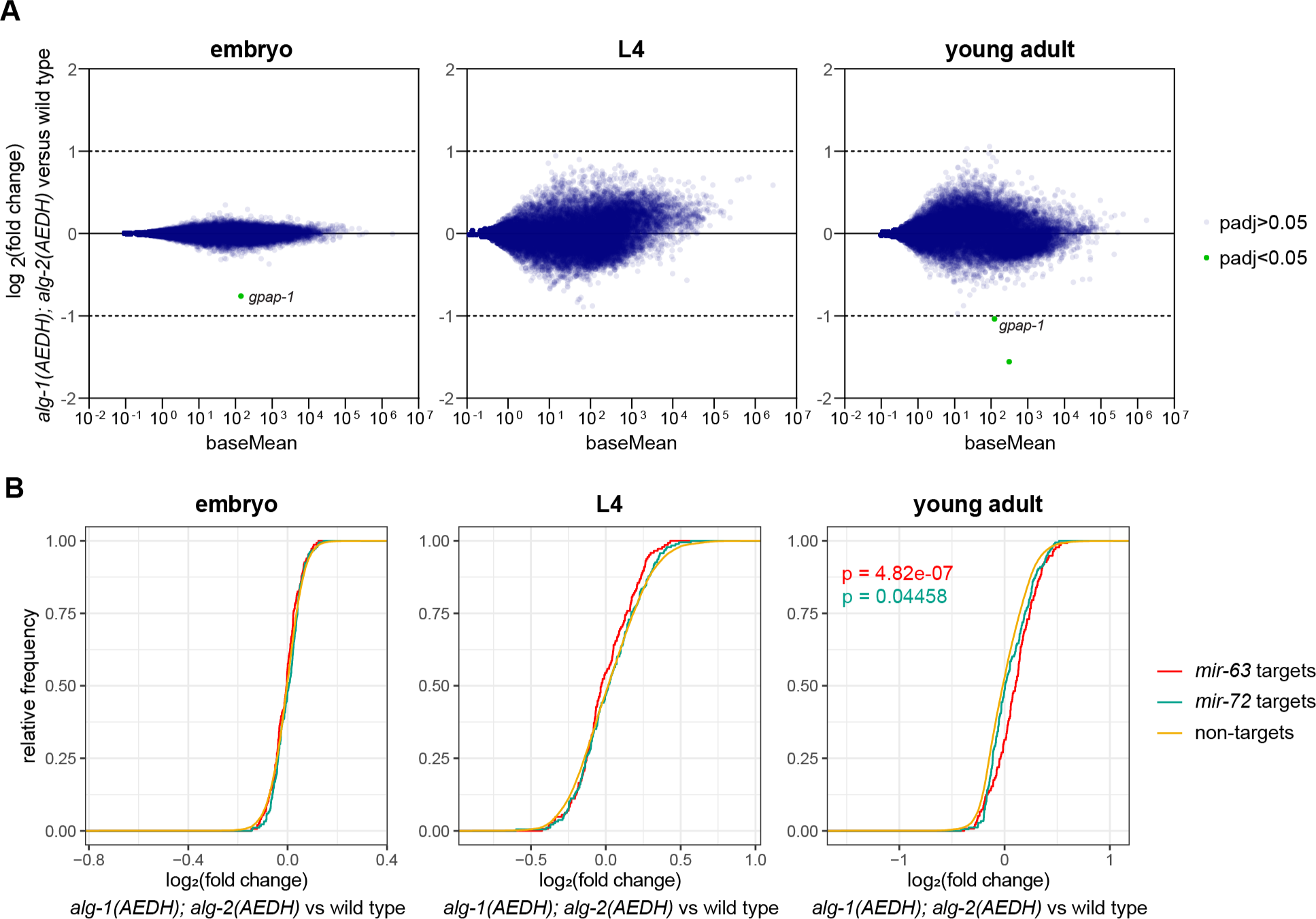
Minor changes in star strand abundance do not result in changes in target repression. A) MA plots showing average abundance (of all genes > 10 reads per million) on X-axis and log2(fold change) on Y-axis in *alg-1(AEDH); alg-2(AEDH)* versus wild type. Only 1, 0, and 2 genes are significantly altered in expression in embryo, L4, and adult, respectively (DESeq2 p_adj_<0.05). B) Empirical cumulative distribution function for log_2_(fold change) values of predicted targets for *mir-63, mir-72* or neither (non-targets). Low abundance transcripts (baseMean < 10) were excluded from the analysis. P-values < 0.05 are shown for Kolmogorov-Smirnoff test (comparison to non-targets). A-B) Three (L4) or four (embryo and young adult) biological replicates of each genotype were analyzed.

To determine whether indirect effects of miRNA star strand stabilization may more subtly impact gene expression in *alg-1(AEDH); alg-2(AEDH)*, we analyzed all Targetscan-predicted targets of *mir-63* and *mir-72* (140). While no significant effect was observed in embryos or L4s, in adults we observed significant upregulation of predicted targets compared to transcripts not targeted by these two miRNAs; the effect was stronger for *mir-63* targets than *mir-72* targets (Figure 5B). This is consistent with partial loss of function of these miRNAs due to failed maturation of a subset of miRISCs in the slicing mutant. Expression of *mir-63* is highest in adult among the stages examined, which may underlie the stage specificity of this effect, and the low guide:star ratio of *mir-63* in *alg-1(AEDH); alg-2(AEDH)* may underlie the larger effect size for *mir-63* than *mir-72* (Figure S6A). Thus, while high-amplitude impacts on individual genes is not observed, the more subtle effects observed by ensemble target analysis are consistent with our model of slicing contributing to star strand unwinding.

We further examined the gene *gpap-1*, which was down-regulated in both embryo and adult *alg-1(AEDH); alg-2(AEDH)* animals (Figure 5A). By close examination of the RNA-seq data, we observed a single-nucleotide polymorphism in the *gpap-1* 5’ UTR in *alg-1(AEDH); alg-2(AEDH)* samples (Figure S6B). Thus, this polymorphism is very likely the basis of these gene expression changes, rather than the mutations in *alg-1* and *alg-2*. No homology was observed between the guide RNAs used to generate the *alg-1* and *alg-2* alleles and the region containing the SNP in *gpap-1*. Thus, we posit that this SNP is due to genetic drift and fixation that occurred during the generation of the *alg-1(AEDH); alg-2(AEDH)* strain. Overall, this suggests that target mRNA cleavage plays little or no role in regulation of gene expression by miRNAs in *C. elegans.* Furthermore, the role that slicing plays in miRISC maturation drives only subtle downstream changes in gene expression.

### Down-regulation of select guide strands in the absence of slicing is *ebax-1-*independent

In addition to elevation of star strands in the *alg-1(AEDH); alg-2(AEDH)* mutants, decreased levels of a few miRNA guide strands were also observed. The three guide strands that were downregulated in two or more developmental stages were *mir-243-3p*, *mir-1829a-5p*, and *mir-1821-3p*. We sought to formulate a model for why these guides might be downregulated in *alg-1(AEDH); alg-2(AEDH)* based on the features of these miRNAs. *mir-243-3p* was previously shown to be largely loaded into RDE-1 and to act as an siRNA to target a fully complementary site in the 3’ UTR of Y47H10A.5 (100). Highly complementary targets can drive target-directed miRNA decay (TDMD); in this process, target cleavage usually does not occur due to a region of central mismatches in the miRNA:target duplex (135, 141–153). A minor portion of *mir-243* is bound by ALG-1, and we wondered whether the mild decrease in *mir-243* (as well as *mir-1829a-5p* and *mir-1821-3p*) in *alg-1(AEDH); alg-2(AEDH)* could be due to enhanced TDMD in the absence of target cleavage (Figure 6A). In this model, TDMD would be less active when ALG-1 is wild type and can cleave the target; when cleavage is inactivated in the *alg-1(AEDH); alg-2(AEDH)* context, TDMD would be more potent, leading to decreased *mir-243* (and other miRNA guide strands) in this mutant (Figure 6A).

**Figure 6.**
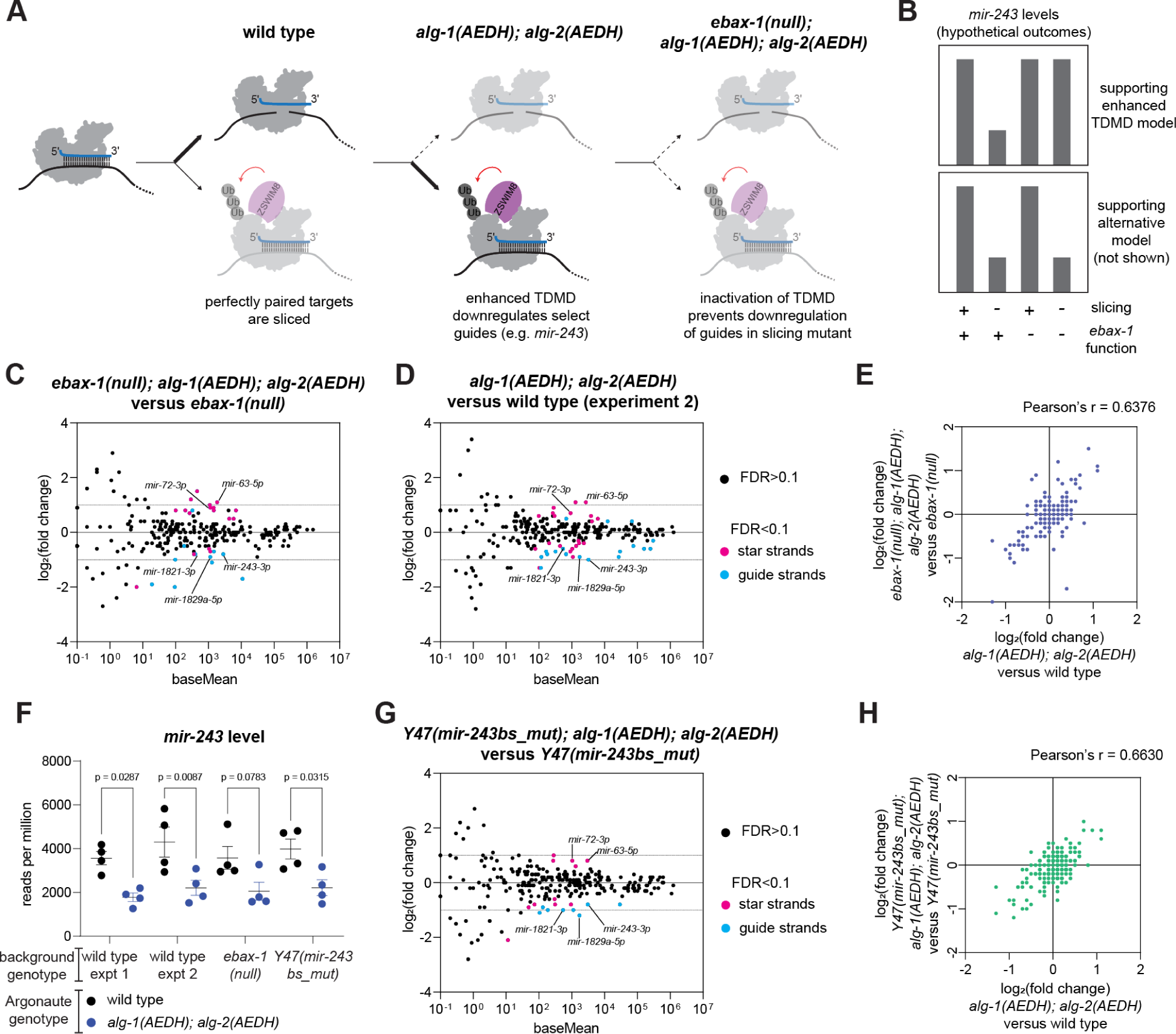
Down-regulation of select guide strands in the absence of slicing is EBAX-1*-* independent. A) Schematic of model being tested. Highly complementary targets that would otherwise be sliced may instead induce TDMD when slicing is inactivated. B) Possible outcomes that would support the model proposed in (A) (top) or support an alternative mechanism (bottom). C-D, G) MA plots showing average abundance on X-axis and log_2_(fold change) on Y-axis. miRNAs showing significant changes are shown in light blue or pink (DESeq2 FDR<0.1). Genotypes being compared are indicated in graph title. E) Correlation of changes induced by *alg-1(AEDH); alg-2(AEDH)* in wild type (X-axis) or *ebax-1(null)* background (Y-axis). F) Level of *mir-243* in each sample after default DESeq2 normalization in pairwise comparisons shown. One-way ANOVA was performed, and p-values are from post hoc Sidak’s multiple comparisons test. E, H) Only miRNAs with baseMean > 50 in wild type experiment 1 are included. (C-H) All samples are young adults. Four biological replicates of each genotype were analyzed.

We tested this model by inactivating this putative TDMD in two ways: 1) by introducing a null mutation in the TDMD effector EBAX-1 (known as ZSWIM8 in mammals) and 2) by deleting the binding site for *mir-243* in the Y47H10A.5 3’ UTR (100, 154–156). If the model is correct, then either of these contexts should abrogate the effect of the *alg-1(AEDH); alg-2(AEDH)* mutations on *mir-243* abundance (Figure 6B). Comparing *ebax-1(null)* young adults to wild type, we corroborate the stabilization of *mir-788, mir-247, mir-42, mir-230, mir-797, mir-43 and mir-253* in this mutant as previously observed (Figure S7, Table S13) (155). (Note that *mir-35-41* are not highly expressed in these samples, which do not contain embryos, in contrast to the previous studies of *ebax-1(null)* which examined gravid adults or embryos (155, 157).)

Next, *ebax-1(null)* was introduced to the *alg-1(AEDH); alg-2(AEDH)* background, and miRNA abundances were compared to *ebax-1(null)* single mutants (156). Very similar amplitude in changes were observed when comparing *ebax-1(null); alg-1(AEDH); alg-2(AEDH)* to *ebax-1(null)* (Figure 6C, Table S14) as when comparing *alg-1(AEDH); alg-2(AEDH)* to wild type (Figure 6D). The correlation of log_2_(fold changes) across all abundant miRNAs in these two comparisons was fairly strong (Figure 6E, Pearson’s r = 0.6376) since it was similar to the correlation between two independent experiments of the same genotypes (each experiment including four biological replicates, Figure S7B, Table S15, Pearson’s r = 0.7173). This demonstrates that overall the observed effects of *alg-1(AEDH); alg-2(AEDH)* are not EBAX-1-dependent.

When examining the guide strands that were decreased in *alg-1(AEDH); alg-2(AEDH)*, similar downregulation was observed for *mir-243-3p*, *mir-1829a-5p*, and *mir-1821-3p* in the *ebax-1(null)* context (Figure 6C, D). Furthermore, the amplitude of downregulation is similar in the *ebax-1(null)* context (Figure 6C, D, F), and our interpretation is that this downregulation is EBAX-1-independent. Thus, the model that this change is due to enhanced TDMD in the catalytic dead mutant is unsupported by the experiments performed.

The *mir-243* binding site mutant (hereafter *Y47(mir-243bs_del)*) further refuted the model. The decrease of *mir-243* was still observed when comparing *Y47(mir-243bs_del); alg-1(AEDH); alg-2(AEDH)* to *Y47(mir-243bs_del)* single mutants (Figure 6F-H, Table S16). This is further evidence negating a role for TDMD in the decrease of select miRNA guide strands in the slicing mutant (also see Discussion).

## Discussion

Here, we demonstrate an unexpectedly modest role for slicing in the miRNA pathway in *C. elegans*. Unlike in vertebrates, no miRNA species are strictly dependent upon slicing for their biogenesis or unwinding. Furthermore, target gene expression is only moderately affected as measured by RNAseq. This results in normal development and reproduction in standard laboratory conditions. This work thus overturns the current literature with regard to the importance of miRNA-guided slicing in *C. elegans* (13). *C. elegans* is the first animal to our knowledge in which slicing by miRNA Argonautes is conserved but dispensable on an organismal level.

As with any genetic study, some form of compensation could ameliorate phenotypes in animals passaged over multiple generations. The catalytic-dead mutations were fairly easy to isolate by CRISPR, and we did not observe stronger phenotypes in early post-CRISPR or post-cross generations, both arguing against strong compensation. Nonetheless, this caveat should be considered, since a compensatory increase in the efficiency of slicing-independent unwinding, for instance, could alleviate the impact of catalytic-dead mutations on RISC maturation.

Another caveat to the interpretation of our results is the lack of validation of the impact of the AEDH mutations on slicing via slicing assays. We attempted to perform these assays using recombinant ALG-1/2 produced in insect cells (Sf9 and Hi5) or *E. coli* [BL21 (RIPL)], and we used the AliCE cell-free protein expression system (158). None of these isolates showed efficient slicing by the wild type proteins. Thus, we were unable to test the impact of the AEDH mutations on slicing, and this remains a caveat to our study, despite extensive characterization of the contributions of the residues of the catalytic tetrad to slicing in the context of other Argonaute proteins (159).

Concurrently to this work, the Simard lab also generated and performed molecular and phenotypic analysis of ALG-1 and ALG-2 slicing mutants (Pal, et al. *Co-submitted*). By examining compound mutants and long-term growth at elevated temperature (25°C), Pal, et al. observed a greater number of de-regulated miRNAs, though largely overlapping with the changes we observed; prolonged growth at high temperature may contribute to these additional expression changes. Pal et al. also observe fitness phenotypes associated with catalytic inactivation of ALG-1 and/or ALG-2 in the context of prolonged high temperature. Thus, although phenotypes of slicing mutants are much milder than previously thought, they are still apparent in sensitized genetic and physical conditions. These stress-associated phenotypes may provide an experimental framework for understanding the selection pressure for maintaining the catalytic activity of ALG-1 and ALG-2 in the long-term.

Molecularly, the basis of these phenotypes may be related to impaired miRNA:star duplex unwinding. We observed a measurable effect on duplex unwinding, represented by a modest overall stabilization of star strands in embryos (Figure 7). Why is this more prominent in *C. elegans* than other previously studied organisms? By examining single mutant strains, catalytic-dead ALG-2 appears to have a greater unwinding defect than catalytic-dead ALG-1. ALG-2 may be more dependent upon slicing for unwinding than previously-examined miRNA Argonautes from other organisms. This may be due to either lower efficiency of slicing-independent unwinding or to higher efficiency of slicing of bulged miRNA:star duplexes. Differences in the N terminal domain or non-catalytic residues in the PIWI domain could contribute to either reduced slicing-independent unwinding or enhanced slicing, whereas differences in the PAZ domain could also impair slicing-independent unwinding (17–20, 51, 80, 81, 160). This opens up new areas of investigation into the biochemical properties of ALG-2 compared to ALG-1 or Argonautes from other species.

**Figure 7.**
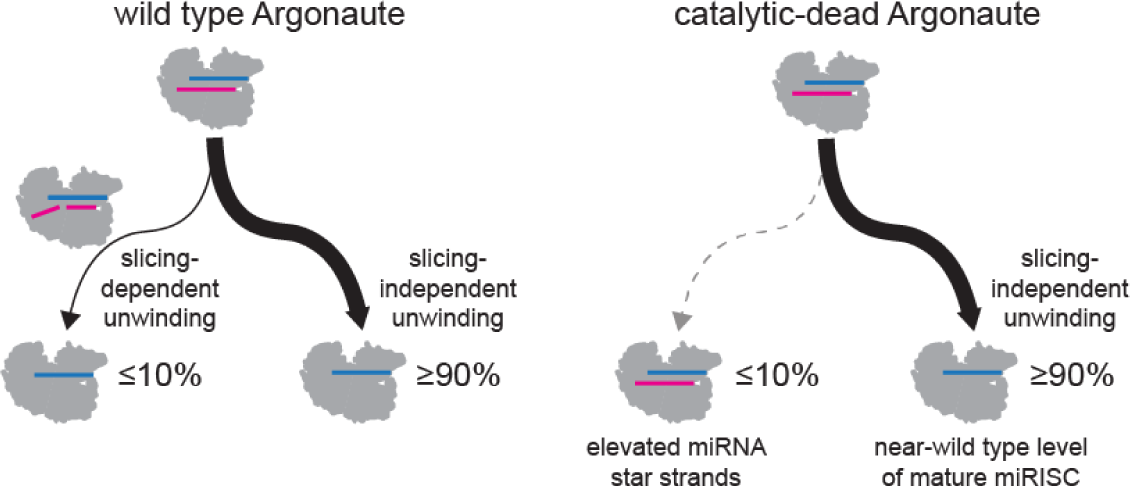
Model of observed effects on miRNA star strands in slicing mutants. A small portion of miRISC is matured by the slicing-dependent unwinding pathway. In *alg-1(AEDH);alg-2(AEDH)* mutant backgrounds, this small portion of miRISC retains the star strand (leading to elevated star strand levels). The major portion of miRISC is matured in a slicing-independent manner, thus leaving overall miRISC function largely intact in the mutant background. Slicing-dependent unwinding is most apparent in embryos in which ALG-2 is highly expressed and for certain thermodynamically-stable miRNA:star duplexes like *mir-63* and *mir-72*. For most miRNAs in L4 and adult samples, slicing-dependent unwinding is negligible.

Some miRNA star strands were elevated in catalytic-dead mutants in all examined contexts, not only in embryos. The duplexes from which these star strands are derived (*mir-63* and *mir-72*) have fairly high thermodynamic stability. For *mir-72*, we find that this duplex stability is indeed a determinant of its dependence on slicing for unwinding. For *mir-63*, lowering duplex stability did not rescue unwinding in the absence of slicing. Thus, duplex stability is *one* of the factors that determines the extent of reliance on slicing for unwinding, as previously observed in other settings (48, 50, 134). However, as illustrated by the case of *mir-63* and the efficient slicing-independent unwinding of even more stable duplexes (Figure 4A), other uncharacterized factors also control the extent of slicing-dependent unwinding. Still, even for *mir-63* and *mir-72,* slicing-dependent unwinding is the minor pathway; only modest elevation in star strands is observed, and the vast majority of miRISC matures to the guide-only bound state and properly represses targets (Figure 7).

A more mysterious observation in this study was the decreased levels of a few guide strands. While we explored a model of enhanced TDMD in the absence of slicing, this was not borne out. However, conventional TDMD has yet to be demonstrated in *C. elegans* (133, 157, 161), so the pathways of slicing and TDMD may not be competing in this context as might be expected in other systems (135, 141–155). Because of the modest reduction of these few guide strands, we posit that slicing is not central to their biogenesis. However, slicing could represent one of multiple biogenesis pathways for these guides or somehow regulate their canonical biogenesis. Interestingly, at least two of the three downregulated guides are primarily loaded into an alternative Argonaute: *mir-243-3p* into RDE-1 and *mir-1829a-5p* into ALG-5 (100, 129). Could ALG-1/2 cleavage products load into other Argonautes? Precedents for this type of interaction include the loading of Aub and Ago3 cleavage products into each other in the piRNA ping-pong pathway (162). The secondary structures of *mir-243* and *mir-1829a* do not provide a clear parallel, but future experiments may nonetheless reveal some similarity.

Overall, this study of miRISC-mediated slicing in a model organism outside the vertebrate lineage showed a surprisingly mild effect on physiology. Although laboratory conditions facilitate the reproducibility to observe clear molecular signatures, they do not recapitulate challenges encountered in the wild. Harsher growth conditions used in a parallel study revealed modest fitness phenotypes associated with loss of miRISC slicing (Pal, et al. *Co-submitted*). The role of slicing in star strand unwinding may suggest a greater role in cold conditions, such as cold shock, where increased duplex stability may disfavor slicing-independent unwinding (50). Robustness of miRISC maturation at a broad range of temperatures may contribute to selection pressure on catalytic residues in the wild. Moreover, our study examined samples derived from whole animals. In the long term, tissue-specific studies may reveal more salient roles for miRISC-mediated slicing not apparent in our study.

## Data Availability

All deep sequencing raw data can be accessed under SRA Bioproject PRJNA922944. Reviewer link: https://dataview.ncbi.nlm.nih.gov/object/PRJNA922944?reviewer=u39pd248iat8khairclnt9ge21

## Acknowledgments

We thank WormBase, the NIDDK Genomics Core, the NCI Genomics Core, and NIH High Performance Computing. Strains are regularly received from and deposited to the CGC, which is funded by NIH Office of Research Infrastructure Programs (P40 OD010440). We thank John Kim for the 3xFLAG::ALG-1 strain (QK67). We thank Amy Pasquinelli and Martin Simard for RNAi clones, and Kevin O’Connell for antibody DM1A. Thank you to Anna Zinovyeva, Martin Simard, and members of the McJunkin lab for helpful discussions.

