## Supplemental Figures for "The catalytic activity of microRNA Argonautes plays a modest role in microRNA star strand destabilization in *C. elegans*"

**List of Supplemental Materials**

**Figure S1.** Western blot of FLAG-tagged wild type and AEDH mutant Argonautes
**Figure S2.** Effect of RNAi on phenotype in AEDH mutant backgrounds
**Figure S3.** Spike in normalization supports minor changes in miRNA abundances
**Figure S4.** Validation of differential expression by qPCR
**Figure S5.** Standard curves for absolute miRNA quantification by qPCR
**Figure S6.** Polymorphism in *gpap-1* 5' UTR likely contributes to decreased expression
**Figure S7.** Reproducibility of miRNA changes in *ebax-1(null)* and across two *alg-1(AEDH); alg-2(AEDH)* experiments

**Table S1.** Strains used in this study
**Table S2.** Alleles generated and oligonucleotides used in this study
**Table S3.** Samples used for deep sequencing
**Table S4.** Raw miRNA and spike-in reads for all samples
**Table S5.** Results of small RNA sequencing analysis in embryo samples*: alg-1(AEDH); alg-2(AEDH)* versus wild type
**Table S6.** Results of small RNA sequencing analysis in L4 samples: *alg-1(AEDH); alg-2(AEDH)* versus wild type
**Table S7.** Results of small RNA sequencing analysis in adult samples: *alg-1(AEDH); alg-2(AEDH)* versus wild type (experiment 1 samples)
**Table S8.** Results of small RNA sequencing analysis in embryo samples: *alg-1(AEDH)* versus wild type
**Table S9.** Results of small RNA sequencing analysis in embryo samples: *alg-2(AEDH)* versus wild type
**Table S10.** Results of RNA-seq analysis in embryo samples: *alg-1(AEDH); alg-2(AEDH)* versus wild type
**Table S11.** Results of RNA-seq analysis in L4 samples: *alg-1(AEDH); alg-2(AEDH)* versus wild type
**Table S12.** Results of RNA-seq analysis in adult samples: *alg-1(AEDH); alg-2(AEDH)* versus wild type
**Table S13.** Results of small RNA sequencing analysis in adult samples: *ebax-1(null)* versus wild type
**Table S14.** Results of small RNA sequencing analysis in adult samples: *ebax-1(null); alg-1(AEDH); alg-2(AEDH) versus ebax-1(null)*
**Table S15.** Results of small RNA sequencing analysis in adult samples: *alg-1(AEDH); alg-2(AEDH)* versus wild type (experiment 2 samples)
**Table S16.** Results of small RNA sequencing analysis in adult samples: *Y47(mir-243bs_del); alg-1(AEDH); alg-2(AEDH)* versus *Y47(mir-243bs_del)*

### Supplemental Figures

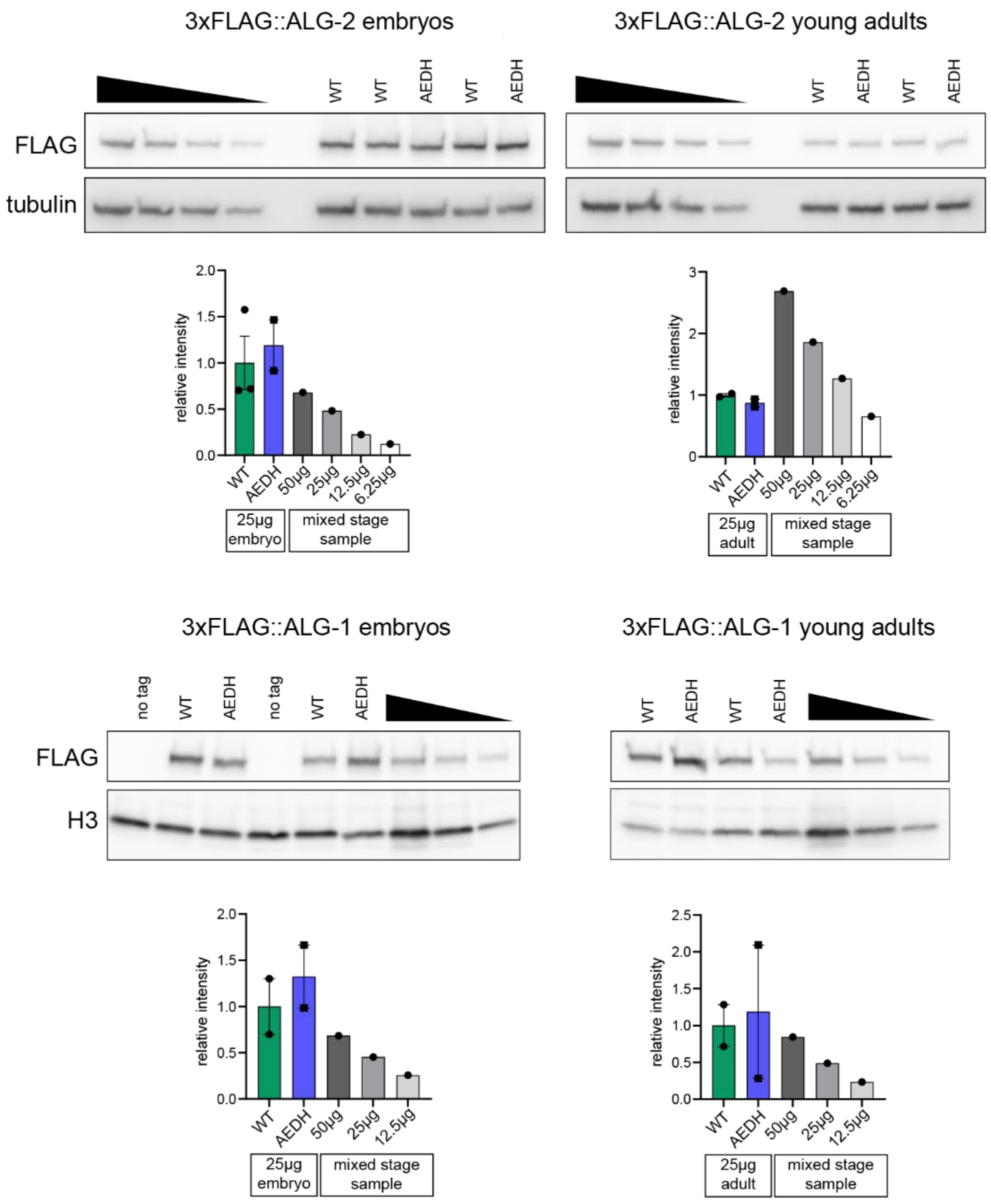


**Figure S1. Western blot of FLAG-tagged wild type and AEDH mutant Argonautes.** Two to three biological replicates of each genotype were quantified. For experimental samples, 25µg of protein is loaded per lane. A dilution series of a mixed-stage sample is included to calibrate fold changes. For experimental samples, FLAG signal was divided by loading control signal in the same lane to determine normalized FLAG signal; for dilution series samples, normalized FLAG signal was calculated by dividing each FLAG signal by the average of the wild type experimental sample loading control signals. Each normalized FLAG signal was then divided by the average normalized FLAG signal for all wild type samples.

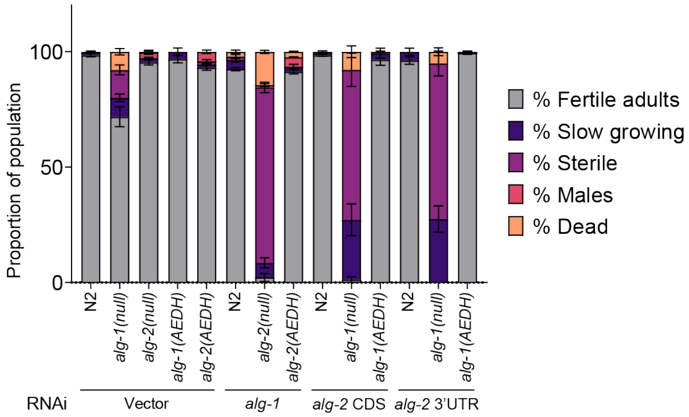


**Figure S2. Effect of RNAi on phenotype in AEDH mutant backgrounds.** RNAi and phenotype scoring was performed as in Bouasker, et al. Eight plates were scored per genotype/RNAi and treated as replicates. n > 470 animals per genotype/RNAi condition. Mean and SEM are plotted. *AEDH* mutations do not behave as loss-of-function alleles since they display no synthetic lethality with RNAi of the other ALG paralog.

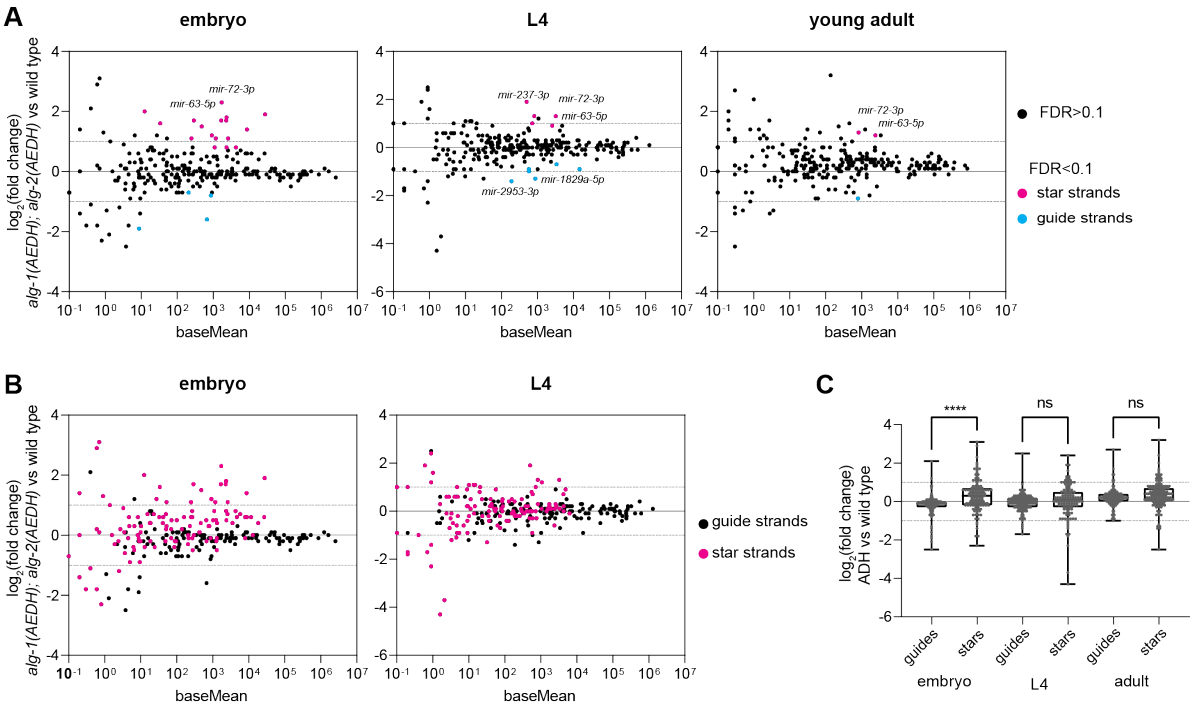


**Figure S3. Spike in normalization supports minor changes in miRNA abundances.** A) MA plot showing average abundance on X-axis and $log_2$(fold change) on Y-axis. miRNAs showing significant changes in *alg-1(AEDH); alg-2(AEDH)* versus wild type are shown in blue (miRNA guide strands) or pink (star strands) (DEseq2 FDR<0.1). Reads were normalized to exogenous spike-ins (See methods and Table S2). Three (L4) or four (embryo and young adult) biological replicates of each genotype were analyzed. (B) MA plot as in (A), with all guide strands shown in black and all star strands in pink. (C) Summary of $log_2$(fold change) values for all guides or star strands in each stage in *alg-1(AEDH); alg-2(AEDH)* versus wild type. $Log_2$(fold change) of star strands were compared to those of guide strands in the same sample by one-way ANOVA followed by Sidak's multiple comparisons test. ***p<0.0001

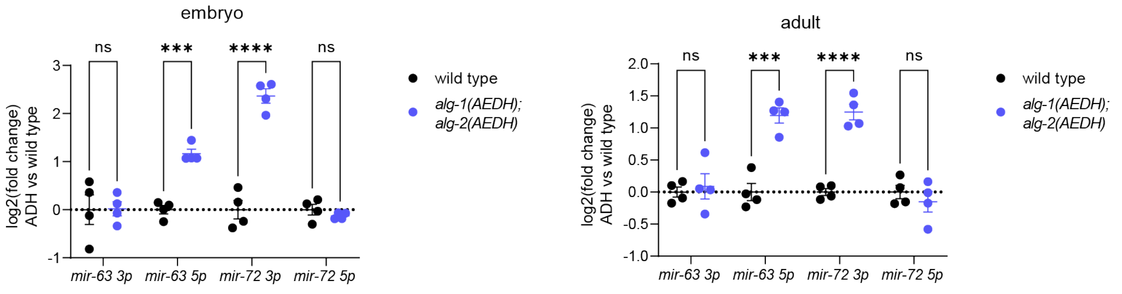


**Figure S4. Validation of differential expression by qPCR.** miRNA Taqman-qRT-PCR assays validate increased levels of *mir-63-5p* and *mir-72-3p* in the *alg-1(AEDH); alg-2(AEDH)* background. Unpaired t-tests, ****p-value < 0.0001, ***p-value < 0.001.

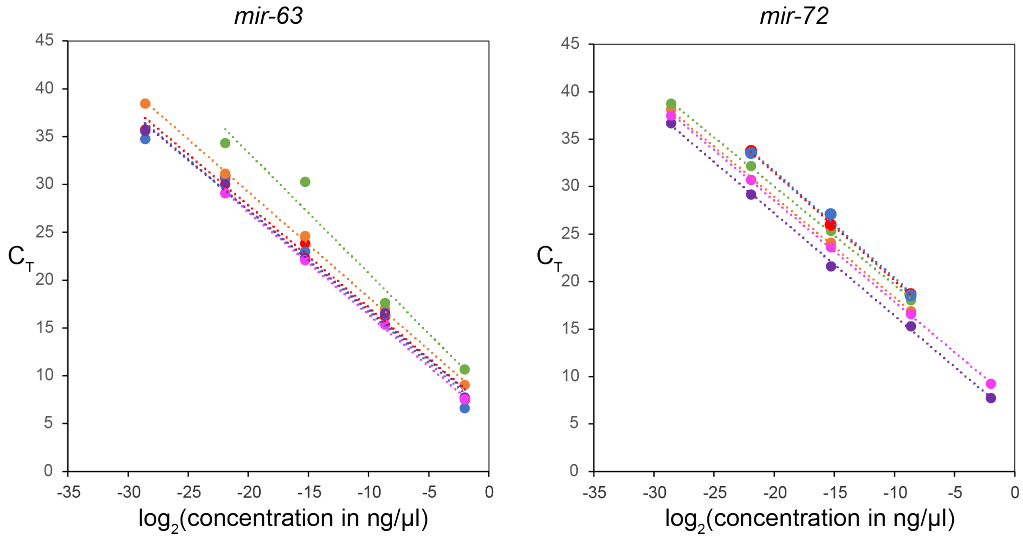


| | | |
|---|---|---|
| mir-63-3p | y = -1.0665x + 6.4959 | $R^2$ = 0.9907 |
| mir-63-5p | y = -1.0633x + 6.0536 | $R^2$ = 0.9816 |
| mir-63-3p_mut2 | y = -1.0993x + 7.2333 | $R^2$ = 0.9988 |
| mir-63-5p_mut2 | y = -1.0436x + 6.4644 | $R^2$ = 0.9944 |
| mir-63-3p_mut3 | y = -1.074x + 5.6509 | $R^2$ = 0.9990 |
| mir-63-5p_mut3 | y = -1.26x + 8.1393 | $R^2$ = 0.9660 |

| | | |
|---|---|---|
| mir-72-3p | y = -1.1342x + 8.7734 | $R^2$ = 0.9998 |
| mir-72-5p | y = -1.1278x + 9.12 | $R^2$ = 0.9933 |
| mir-72-3p_mut1 | y = -1.0592x + 7.7451 | $R^2$ = 0.9996 |
| mir-72-5p_mut1 | y = -1.0799x + 5.5879 | $R^2$ = 0.9992 |
| mir-72-3p_mut2 | y = -1.0636x + 7.2479 | $R^2$ = 0.9998 |
| mir-72-5p_mut2 | y = -1.0377x + 9.2635 | $R^2$ = 0.9995 |

**Figure S5. Standard curves for absolute miRNA quantification by qPCR.** Synthetic RNA oligonucleotides were diluted to indicated concentrations, and 1.66µl was used in 5µl RT reactions and downstream miRNA-Taqman as indicated in methods. Average of three technical replicates is shown. Highest and lowest concentrations were excluded from the curve if they were outside the linear range. All experimental sample concentrations fell within the linear range of the respective assay.

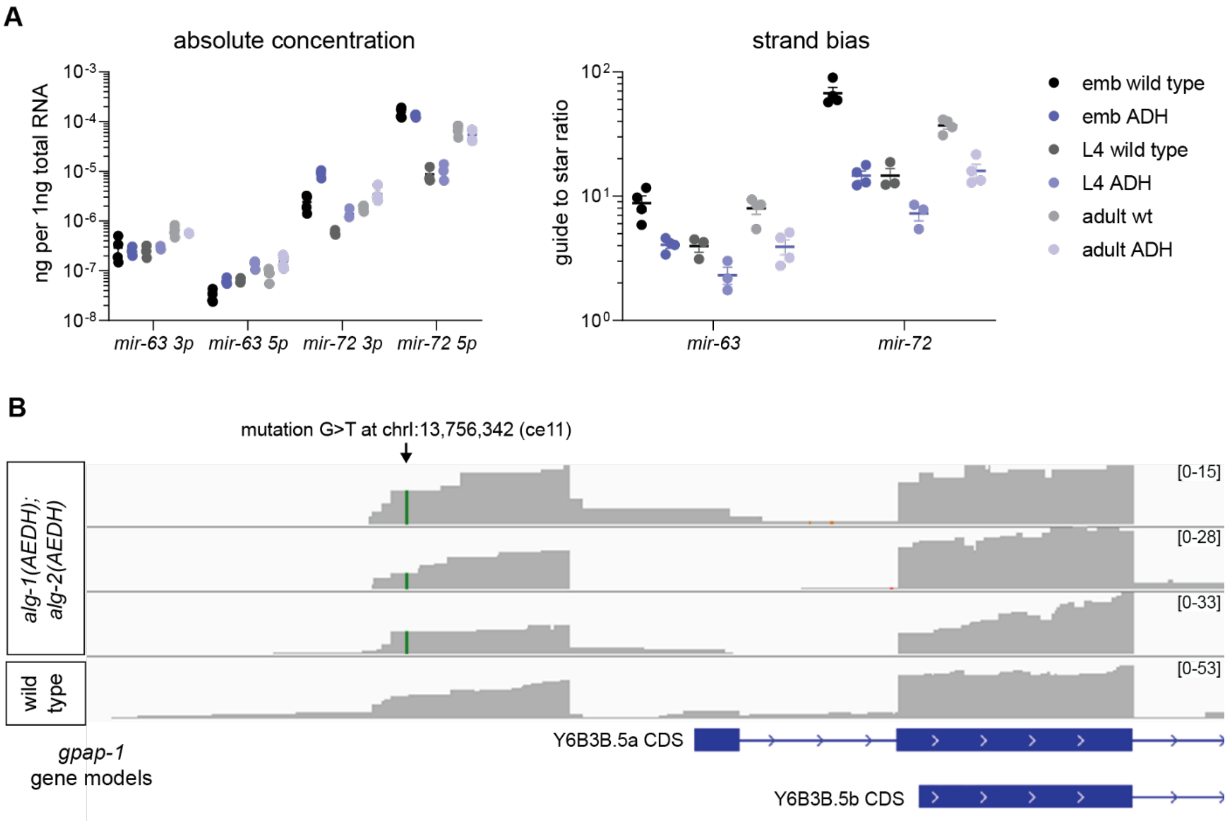


**Figure S6. Polymorphism in *gpap-1* 5' UTR likely contributes to decreased expression.** A) Left: Absolute quantification by qPCR of *mir-63* and *mir-72* guide and star strands Right: Strand bias calculated from absolute quantification by qPCR. Mean and SEM of three to four biological replicates per condition are shown. B) One gene was identified as mis-regulated across more than one developmental stage in the *alg-1(AEDH);alg-2(AEDH)* mutant strain. Close examination of RNA-seq data revealed that this gene, *gpap-1*, contains a polymorphism in its 5' UTR in the mutant strain.

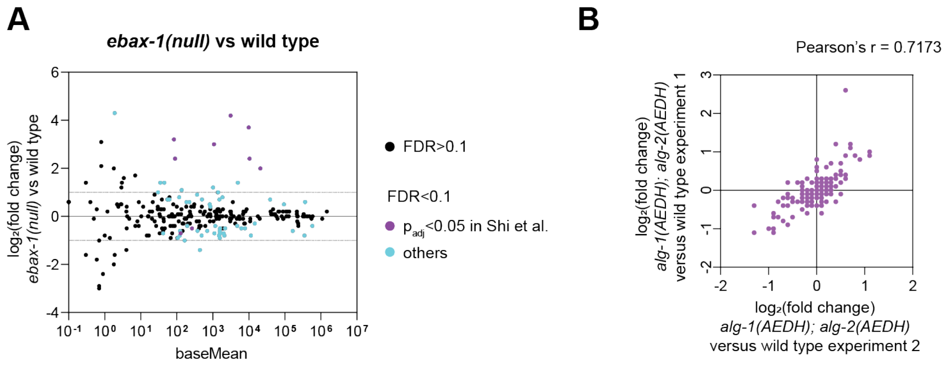


**Figure S7. Reproducibility of miRNA changes in *ebax-1(null)* and across two *alg-1(AEDH); alg-2(AEDH)* experiments.** A) MA plots showing average abundance on X-axis and $\log_2$(fold change) on Y-axis. miRNAs showing significant changes are highlighted in light blue or purple (DEseq2 FDR<0.1); upregulated miRNAs that were previously noted in Shi, et al are in purple. Four biological replicates of each genotype were analyzed. B) Correlation of changes induced by *alg-1(AEDH); alg-2(AEDH)* in wild type experiment 1 (X-axis) or experiment 2 (Y-axis). Each experiment consists of four biological replicates of each genotype. Only miRNAs with baseMean > 50 in wild type experiment 1 are included. (A-B) All samples are from young adults.
